# Investigating the role of RNA-binding protein Ssd1 in aneuploidy tolerance through network analysis

**DOI:** 10.1101/2024.07.19.604323

**Authors:** H. Auguste Dutcher, Audrey P. Gasch

## Abstract

RNA-binding proteins (RBPs) play critical cellular roles by mediating various stages of RNA life cycles. Ssd1, an RBP with pleiotropic effects, has been implicated in aneuploidy tolerance in *Saccharomyces cerevisiae* but its mechanistic role remains unclear. Here we used a network-based approach to inform on Ssd1’s role in aneuploidy tolerance, by identifying and experimentally perturbing a network of RBPs that share mRNA targets with Ssd1. We identified RBPs whose bound mRNA targets significantly overlap with Ssd1 targets. For 14 identified RBPs, we then used a genetic approach to generate all combinations of genotypes for euploid and aneuploid yeast with an extra copy of chromosome XII, with and without *SSD1* and/or the RBP of interest. Deletion of 10 RBPs either exacerbated or alleviated the sensitivity of wild-type and/or *ssd1*Δ cells to chromosome XII duplication, in several cases indicating genetic interactions with *SSD1* in the context of aneuploidy. We integrated these findings with results from a global over-expression screen that identified genes whose duplication complements *ssd1*Δ aneuploid sensitivity. The resulting network points to a sub-group of proteins with shared roles in translational repression and p-body formation, implicating these functions in aneuploidy tolerance. Our results reveal a role for new RBPs in aneuploidy tolerance and support a model in which Ssd1 mitigates translation-related stresses in aneuploid cells.

## INTRODUCTION

RNA-binding proteins (RBPs) are critical mediators of cellular responses, with highly interconnected functions often spanning multiple stages of RNA life cycles (Müller-Mcnicoll and Neugebauer 2013; Goswami et al. 2024; Hogan et al. 2008). Many RBPs are both complex in sequence—with many functional domains—and pleiotropic in either functions or phenotypes they affect (Hentze et al. 2018; Corley et al. 2020). As a group they are integral to all facets of post-transcriptional regulation, including but not limited to RNA transcription, nuclear export, processing, subcellular localization, translation, stability, and decay (Dreyfuss et al. 2002; Mazumder et al. 2003). It is not uncommon for RBPs to operate across more than one of these regulatory realms. The pleiotropic nature of RBP functions often presents a challenge in elucidating their biochemical mechanisms and how those mechanisms impact cellular phenotypes.

Previous work in our lab found that RBP called Ssd1 enables budding yeast to tolerate extra chromosomes, a state known as aneuploidy (Hose et al. 2020; Rojas et al. 2024). Despite the clear implication of Ssd1 in aneuploidy tolerance, the mechanistic basis for its involvement has remained unknown. Many different studies identified Ssd1 in suppressor screens, linking it to processes ranging from cell wall integrity and thermotolerance to tRNA modifications, aging, quiescence, and now aneuploidy (Kaeberlein and Guarente 2002; Xu et al. 2019; Miles et al. 2019; Hu et al. 2018; Li et al. 2009; Hose et al. 2020). Ssd1 is important for delivery of cell wall-associated mRNAs to sites of polarized growth under normal conditions or to RNA-protein granules called p-bodies (PB) during stress (Kurischko, Kim, et al. 2011; Kurischko, Kuravi, et al. 2011). In fact, Ssd1 is a component of PB, although found in only a subset of PBs in the cell (Xing et al. 2020). Ssd1 has also been implicated in translational repression, based on global polysome profiles as well impacts on protein abundance from specific directly bound transcripts (Wanless et al. 2014; J. M. Jansen et al. 2009; Hu et al. 2018; Hose et al. 2020). Unlike its mammalian ortholog Dis3L2 that degrades poly-uridylated transcripts, Ssd1 is catalytically inactive but directly binds several hundred mRNAs at sequences primarily in the 5’ UTR and coding regions (Hose et al. 2020; Bayne et al. 2022; Hogan et al. 2008; J. M. Jansen et al. 2009; Ohyama et al. 2010; Wanless et al. 2014). The RNAs bound by Ssd1 are enriched for those encoding specific functional groups, including cell cycle regulation, cell wall integrity, RNA metabolism, budding, and sterol transport, but they also include many mRNAs outside of these functional groups (Hose et al. 2020; Hogan et al. 2008; Bayne et al. 2022). Although Ssd1 has been linked to mRNA localization and translational control (Wanless et al. 2014; J. M. Jansen et al. 2009; Hogan et al. 2008; Kurischko, Kuravi, et al. 2011; Xu et al. 2019; Hu et al. 2018; Khonsari et al. 2021), how it acts mechanistically is not known. Furthermore, it is unclear which of its functions, bound transcripts, or their related processes influence aneuploidy sensitivity.

One strategy to pinpoint which of Ssd1’s functions are important is to explore other RBPs that may work with Ssd1 by binding the same transcripts. Previous studies in *Saccharomyces* cerevisiae and humans revealed that subsets of RBPs show substantial overlap in mRNA targets (Hogan et al. 2008; Achsel and Bagni 2016; Kershaw et al. 2023). This interconnectivity is an asset to elucidate RBP networks, which can be constructed based on the overlap in RBP targets and on physical interactions between RBPs. The resulting networks can reflect on regulatory principles and shared regulation across specific mRNAs. For example, yeast mRNAs can be effectively categorized into discrete modules based on the RBPs that bind them (Costello et al. 2015; Kershaw et al. 2023). Features of translational control can be inferred by relating these regulatory modules to biophysical properties of the affected mRNAs (Costello et al. 2015; Kershaw et al. 2023). Thus, a network perspective provides greater biological insights into the regulation of mRNA targets than if their interactions were examined piecemeal.

Here we used a network-based approach to better understand the role of Ssd1 in aneuploidy tolerance. Under the hypothesis that Ssd1 may function in this role with other proteins that bind the same RNAs, we first identified a network of RBPs whose bound mRNAs overlap with Ssd1’s known targets. We then experimentally perturbed this network by deleting RBPs in euploid and aneuploid cells, alone and in a *ssd1Δ* background. This allowed us to investigate the influence of each RBP on aneuploidy tolerance, alone and in the context of *SSD1* deletion. We integrated these findings with recent results from a global gene-duplication screen to further refine hypotheses about Ssd1’s functions in aneuploidy tolerance across chromosome duplications. Our results implicate a subnetwork of RBPs and translation factors whose deletion or over-expression influences aneuploidy tolerance in yeast. This work adds to a growing body of evidence that Ssd1 and it is broader RBP network mediate aneuploidy tolerance through functions related to translational suppression. We discuss models for how translational repression could influence tolerance of extra chromosomes.

## RESULTS

### Ssd1 is part of a highly interconnected RBP network

We previously used RNA-immunoprecipitation and sequencing (RIP-seq) to identify 286 mRNAs bound by Ssd1 (Hose et al. 2020). To identify other RBPs that might act with Ssd1 including in mediating aneuploidy tolerance, we identified RBPs that bind more Ssd1 targets than expected by chance. We first collated from public datasets available at the time the mRNAs bound by 65 different RBPs that share at least one Ssd1 target (see Methods and Supplemental Data Table S2). We then identified the subset of RBPs whose bound mRNAs statistically significantly overlap Ssd1-bound mRNAs (FDR < 0.05, hypergeometric test, see Methods).

This approach identified 14 RBPs as part of the Ssd1 network (Figure 1). Several of these proteins are implicated in multiple processes related to RNA biogenesis, and together, the network spans functions related to most stages of mRNA life cycles, including transcript maturation and nuclear export (Isw1 and Gbp2), subcellular localization (She2), translational regulation (Gis2, Mpt5, Puf6, Whi3, and Hek2, among others) and roles in mRNA decay and stability (Ccr4, Dhh1, Pat1, Pub1, Mrn1, Nab6, and Whi3). Like Ssd1, many of these RBPs were previously found to bind cell wall-related transcripts (She2, Hek2, Gbp2, Pub1, Mpt5, Mrn1, and Nab6) or have been otherwise implicated in cell wall integrity (Mpt5, Ccr4, and Hek2) (Kaeberlein and Guarente 2002; Hogan et al. 2008; Ito et al. 2011; Hall and Wallace 2022; Bresson et al. 2023). However, overlap with Ssd1 targets remained significant (FDR < 0.05) for all of these proteins except Pub1 and Nab6 when transcripts corresponding to the GO term “cellular wall” were removed, suggesting that Ssd1’s network is not restricted to cell wall transcript-binding RBPs alone. Interestingly, in addition to sharing mRNA targets with Ssd1, many of these RBPs interact physically and/or genetically with Ssd1 or other RBPs in the network. Ccr4 in particular, along with Ssd1 itself, exhibits many genetic interactions with other RBPs in the network. Furthermore, 11 of the 14 proteins can localize to membraneless condensates including PB or stress granules (SG) that form after stress (reviewed in Escalante and Gasch 2021). Indeed, Ssd1 is known to localize to a subset of PB and SG during various stresses (Kurischko, Kim, et al. 2011; Xing et al. 2020). Together, these results suggest that Ssd1 and its targets are part of a highly interconnected network of proteins that act on similar targets.

**Figure 1.**
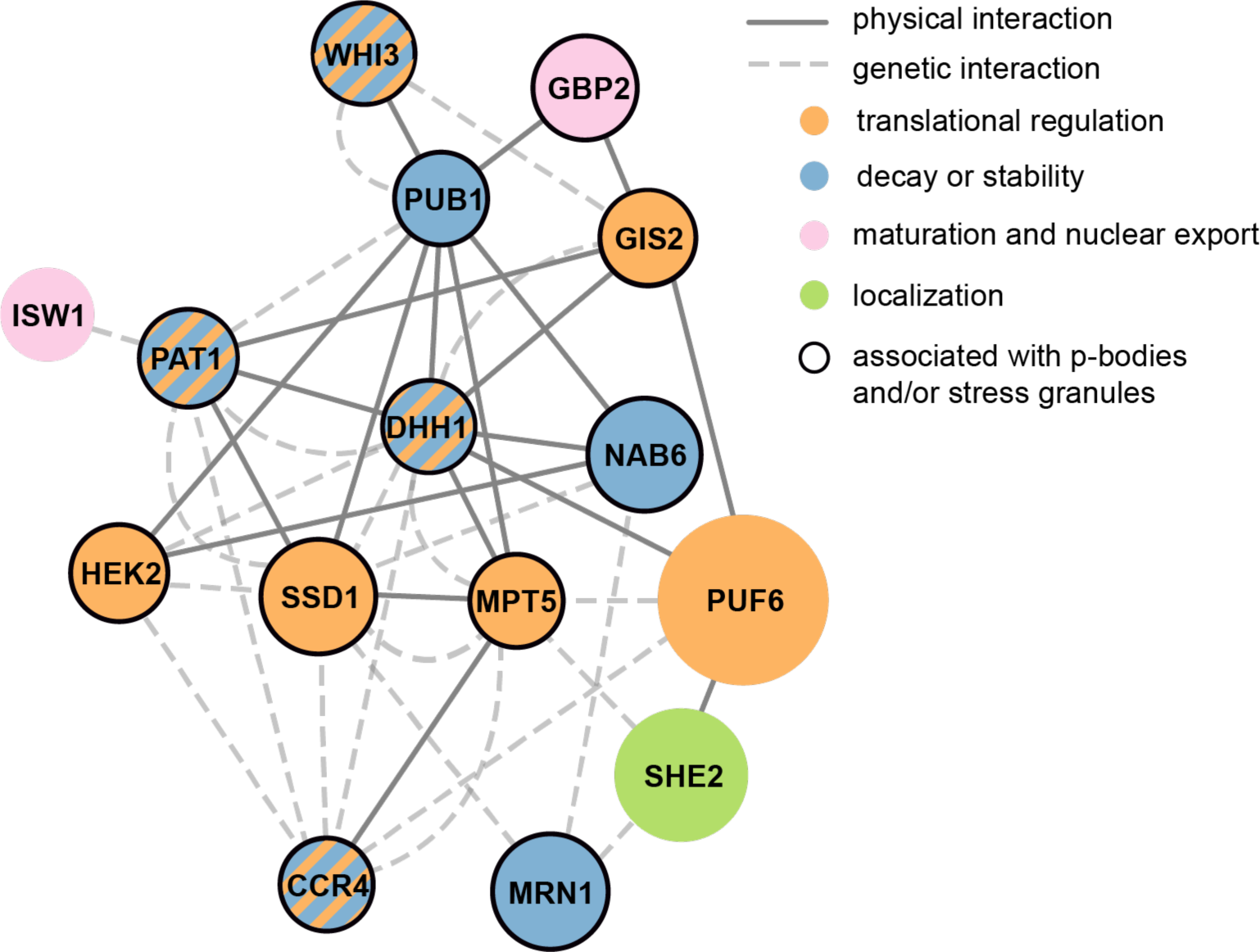
Ssd1 shares a regulatory network with 14 other RNA-binding proteins. Each circle represents an RBP and each line represents a physical or genetic interaction according to the key. With the exception of Ssd1, nodes are scaled to reflect the proportion of that RBP’s targets that are also bound by Ssd1. Colors depict primary functional groups of each RBP as shown; since many RBPs are involved in multiple functions, this figure depicts simplified annotations.

### Many of the RBPs in Ssd1’s regulatory network exhibit synthetic genetic interactions with *ssd1Δ* and aneuploidy

We were especially interested if any of these RBPs, like Ssd1, play a role in aneuploidy tolerance. To investigate this, we devised a genetic strategy to sensitively measure the interaction of each RBP deletion with *SSD1* deletion and chromosome XII (Chr12) duplication, alone and in combination. We chose this chromosome duplication because it is a good representative in our strain background and was available at the time of our analysis. Like in many other aneuploid strains, Chr12 duplication is tolerated in wild-type cells but very deleterious in an *ssd1*Δ background (Rojas et al. 2024). At least part of the dependence on Ssd1 is due to the amplification of specific mRNAs on Chr12 that are very deleterious in a euploid strain lacking *SSD1* (Dutcher et al. 2024).

We engineered a diploid parent in our wild strain background, oak-soil strain YPS1009, that is trisomic for Chr12 (“3nChr12”) and hemizygous at both the *SSD1* locus and the locus of each RBP of interest. We then dissected meiotic products to isolate discrete genotypes that all grow side-by-side in the same environment (Figure 2A). Colony size was determined after a defined growth period and used as a proxy for growth rate. Each colony’s genotype was subsequently determined using drug markers to identify *SSD1* and RBP knockouts, and euploid or aneuploid status was identified by flow cytometry to infer DNA content (Chen et al. 2012). Progeny were diploid due to mating-type switching in this strain background, with the exception of *she2Δ* and *whi3Δ*. These two mutants harbored DNA content consistent with mixed ploidy, likely caused by impaired mating or mating-type switching (R. P. Jansen et al. 1996; Nash et al. 2001), and were thus reconstructed as haploids for further analysis. We compared colony sizes of all genotypes from a single parental strain grown in parallel under identical conditions, including strains with and without *SSD1*, each RBP of interest, an extra copy of Chr12 (hereafter referred to simply as “aneuploidy”), and every combination thereof. Statistically significant genetic interactions were identified if the measured colony size in the double mutant was more or less than expected based on independent action of each RBP and of aneuploidy (through a multiplicative model, see Methods). This method implicated RBP deletions that alleviate or exacerbate aneuploidy stress in wild-type cells and in the context of *ssd1Δ* (see Methods).

**Figure 2.**
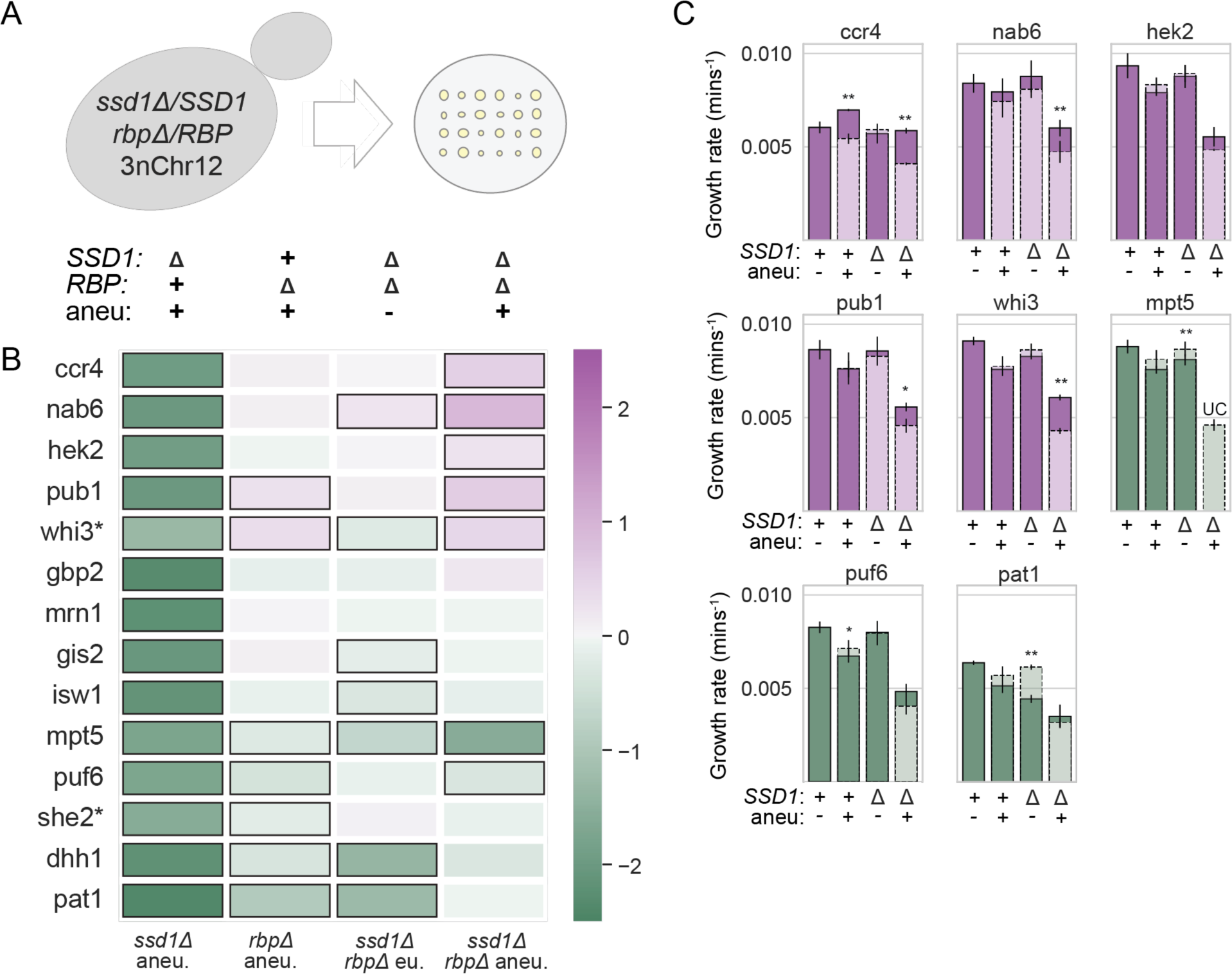
RNA-binding proteins with shared targets exhibit synthetic genetic interactions with *ssd1Δ* and aneuploidy. (A) Dissection of a parent strain trisomic for chromosome 12 (3nChr12) and hemizygous at *SSD1* and each RBP locus of interest yields colonies with all possible genotypes, a subset of which are represented in B. (B) The log_2_(fold-difference) in colony size comparing mean observed size vs mean expected size across batches (see Methods), according to the color scale, for genotypes indicated along the top in (A). Black outlined boxes identify significant effects (FDR <= 0.05). * indicates haploids. (C) Average and standard deviation of exponential growth rates of the denoted strain under standard conditions; all strains shown for each mutant lack the RBP in question. Measured growth rates are represented as solid bars; expected growth rates are represented by light colors and dashed lines; bars with solid color above the dashed line grew better than expected. Mutants with primarily a positive (alleviating) effect are shown in purple; mutants that mostly exacerbated defects are shown in green. (**) p-value <= 0.01, (*) p <= 0.05, (+) p <= 0.1, two-tailed, replicate-paired t-test. UC = unculturable strain.

Our strategy revealed a remarkably high hit rate, with 10 of the 14 RBPs having a significant impact aneuploidy and/or aneuploidy in the context of *ssd1Δ* (Figure 2B). The approach confirmed the expected interaction between *ssd1Δ* and Chr12 duplication in every single dissection, recapitulating the very strong negative genetic interaction between *ssd1Δ* and aneuploidy (see Figure 2B, column 1). No other RBP deletion sensitized cells to chromosome amplification to nearly the same extent as *ssd1Δ*; however, *mpt5Δ, puf6Δ, she2Δ, dhh1Δ,* and *pat1Δ* all sensitized cells to aneuploidy to some degree in wild-type cells (Figure 2B, column 2, FDR < 0.05). Of these, a subset (*mpt5Δ, dhh1Δ,* and *pat1Δ*) also had negative interactions with *ssd1Δ* in a euploid context (Figure 2B, column 3), consistent with previous reports (Moriya and Isono 1999; Costanzo et al. 2016; Kaeberlein and Guarente 2002). Only *mpt5Δ* and *puf6Δ* had significant negative interactions with *ssd1Δ* in an aneuploid background, although the effect of *dhh1Δ* just missed the significance cutoff (Figure 2B, column 4). Thus, several mutants displayed negative genetic interactions with aneuploidy, with *ssd1Δ*, or both, such that they exacerbate the sensitivity to extra Chr12.

Interestingly, a large group of significant effects comprised RBPs whose deletion resulted in larger than expected colonies for aneuploid double-mutants (*ssd1Δ rbpΔ*, Figure 2B column 4). Deletion of *CCR4, NAB6, HEK2, PUB1, or WHI3* in combination with *ssd1Δ* and Chr12 duplication all produced colonies larger than expected from a multiplicative model (see Methods), to varying degrees. In all cases, this positive genetic interaction was limited to or stronger in the aneuploid double mutant, producing a larger effect than seen in the euploid double mutant. Thus, deletion of over a third of the queried RBPs in Ssd1’s network produce significant, reproducible differences in colony size that are specific to the context of *ssd1Δ* and aneuploidy.

One possibility is that cell size and/or colony morphology, rather than growth rate, explain the observed differences in colony size. To measure effects that are truly due to growth rate differences, we measured the growth rate of select RBP deletions in liquid culture, this time in a haploid background to identify ploidy-independent effects (Figure 2C). All 5 positive genetic interactions between RBP deletion, *ssd1Δ*, and aneuploidy were validated in liquid culture (although the *ssd1Δ hek2Δ* aneuploid missed the cutoff for significance). (We noted substantial colony size variability in streaks of *whi3Δ* and *ssd1Δ whi3Δ* strains, which could suggest populations of mixed ploidy, consistent with reports of increased rates of ploidy increase in *whi3Δ* mutants from a previous study (Schladebeck and Mösch 2013)). Growth rate measurements likewise validated most negative genetic interactions between *ssd1Δ* and aneuploidy: in the most striking example, *ssd1Δ mpt5Δ* aneuploids proved entirely unculturable, consistent with the dramatic growth defect observed from our colony size assay. Interestingly, *puf6Δ* and *pat1Δ* liquid growth rates produced mixed results: both deletions exacerbated the sensitivity of *SSD1+* cells to Chr12 duplication, but slightly alleviated the growth defect of *ssd1Δ* aneuploid cells (though missed the cutoff for significance). In summary, most phenotypes from our colony-size assay were validated in liquid growth, confirming that the genetic interactions affect growth rate. Thus, we identified several new proteins in Ssd1’s network that also exhibit intriguing genetic interactions with aneuploidy.

### Incorporating beneficial gene duplications expands the Ssd1-aneuploidy network

Through other work, we recently identified gene duplications that alleviate the growth defect of *ssd1Δ* aneuploids carrying different chromosome amplifications, including Chr12. Genes that benefited multiple *ssd1Δ* aneuploids were enriched for genes involved in RNA metabolism and translation (Dutcher et al. 2024). To further refine hypotheses about Ssd1’s role in aneuploidy tolerance, here we integrated those results with the RBP network presented here. We started with 316 gene duplicates that specifically benefit the *ssd1Δ* Chr12 aneuploid strain (FDR < 0.05, see Dutcher et al. 2024). This group represented diverse functions but was enriched for proteins involved in mRNA processing (p = 7.2E-05, hypergeometric test) and regulation of translation (p = 8.2E-03). We identified the subset of 66 genes that belonged to these and related functional groups (namely, rRNA processing and ribosome biogenesis, splicing, and tRNA-related, see Methods) and combined them into a joint network with RBPs identified above. We considered gene duplicates beneficial to *ssd1Δ* cells with a Chr12 duplication as well as the subset of those genes that also benefic other *ssd1Δ* aneuploids.

The resulting network suggested functional connections among beneficial gene duplicates and impactful RBPs (Figure 3). First, the network clarified functions of beneficial genes: most of the genes related to rRNA, tRNA, and ribosome biogenesis were only scored as beneficial upon duplication of Chr12, which contains the rDNA locus. This suggests that some of the genes may complement Ssd1-dependent effects related to ribosome biogenesis or function. In contrast, duplication of other genes, including those linked to splicing, mRNA processing, and especially translation, were beneficial to multiple chromosome duplications, strongly suggesting more generalizable effects in *ssd1Δ* aneuploids. Interestingly, several of the proteins encoded by beneficial gene duplicates interact physically with impactful RBPs (Figure 3, orange edges). Several interacted with more RBPs in our original network than expected by chance (based on their total number of protein interactions, FDR < 0.05, hypergeometric test, Figure 3, large nodes). Many of these are translational regulators. In fact, the group of 15 beneficial genes that interact with more RBPs than expected was statistically significantly enriched for translational regulators (p = 2.4E-05, hypergeometric test). These enrichments further suggest that Ssd1’s role in aneuploidy tolerance may be linked translation (see Discussion).

**Figure 3.**
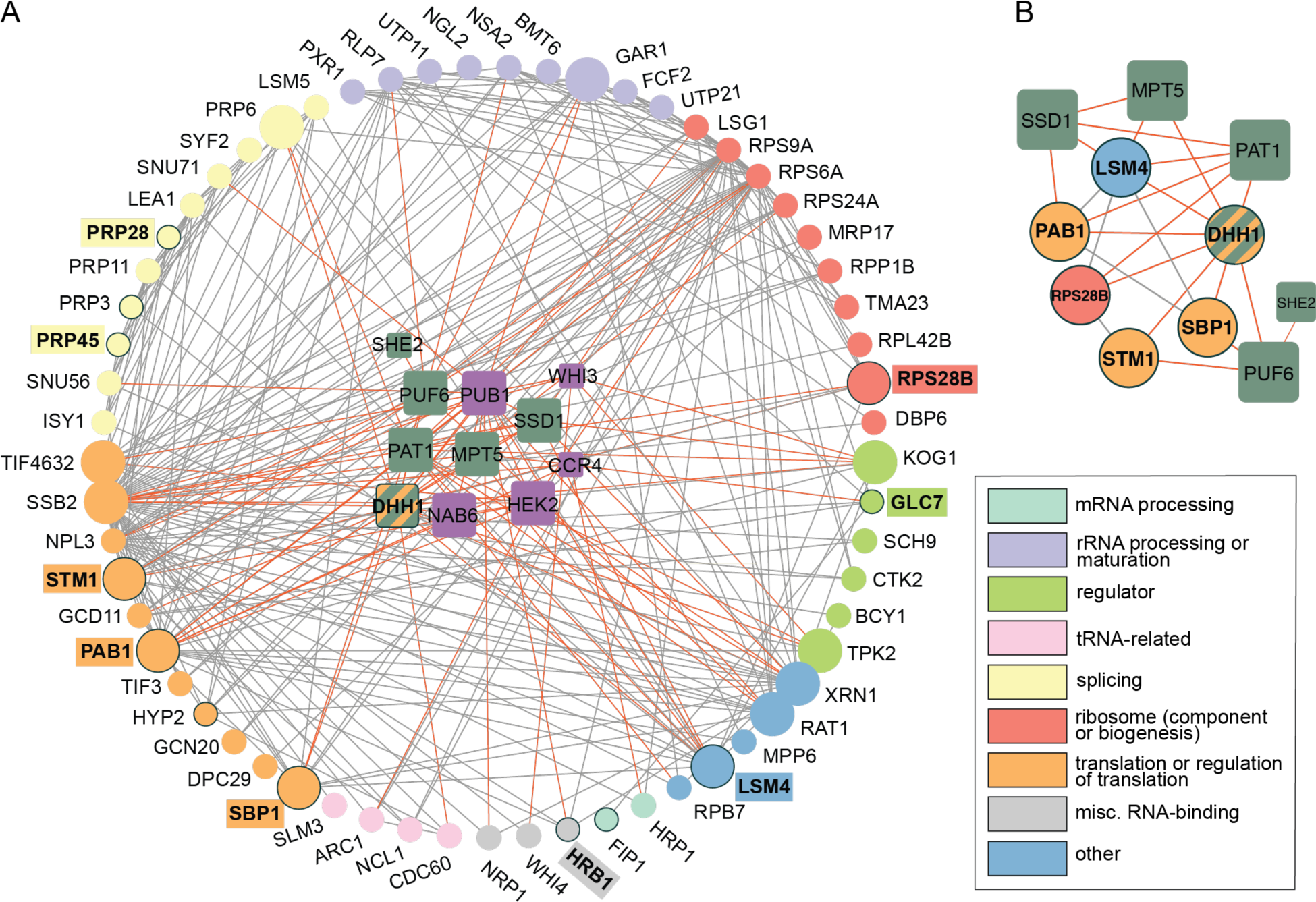
RBPs that genetically interact with aneuploidy physically interact with duplicated proteins that complement aneuploid *ssd1Δ* defects. (A) 65 peripheral nodes (circles) represent genes related to translation and RNA biology (colored with simplified annotations according to the legend) whose duplication is especially beneficial to Chr12 *ssd1Δ* aneuploids. Central nodes (squares) represent significant RBPs from Figure 2, colored purple or green if the deletion alleviates or exacerbates *ssd1Δ* Chr12 sensitivity. Nodes with a black border (label with colored background) indicate gene duplications that confer a benefit in multiple *ssd1Δ* aneuploids carrying different chromosome amplifications. Edges represent physical interactions between encoded peripheral proteins (grey) or with an RBP (orange). (B) Complementary network of gene duplicates beneficial to multiple *ssd1Δ* aneuploids (circles) and RBP deletions (squares) that exacerbate the aneuploid growth defect. *DHH1* belongs to both groups.

To further investigate, we generated a subnetwork of genes linked to complementary impacts on *ssd1Δ* aneuploidy: a benefit when over-produced and/or a defect when deleted. We reasoned that the complementary effects of their genetic perturbation would tap into a network of gene products that function together in the same required process. We took six proteins that are beneficial to multiple *ssd1Δ* aneuploid strains (Dutcher et al. 2024) and show more physical interactions with RBPs studied here (FDR < 0.05). We then combined this list with the six RBPs whose deletion exacerbates growth defects in the presence of Chr12 duplication (Figure 2B). The DEADBOX RNA helicase Dhh1 was identified in both studies: its duplication alleviates the *ssd1Δ* aneuploidy growth defects whereas its deletion sensitizes cells to Chr12 duplication.

The resulting subnetwork implicated deep connections between RBPs (all of which share targets with Ssd1) and genes whose duplication benefits aneuploidy tolerance in the absence of *SSD1*. Remarkably, nearly all of these proteins can be related to translational control. This includes several proteins implicated in translation or ribosome functions (Pab1, Rps28b, Stm1), translational repression specifically (Sbp1, She2 and Puf6, Mpt5, Pab1, Dhh1, along with Ssd1), or RNA decay (Lsm4, Pat1, Dhh1). Strikingly, most of these proteins either localize to PB or influence their function (Pat, Dhh1, Lsm4, Mpt5, Ssd1, Rps28b, and others). Together, our computational and experimental analysis support a role for translational control in mediating aneuploidy tolerance. Implications and models are discussed below.

## DISCUSSION

While Ssd1 is clearly involved in aneuploidy tolerance, its precise function in mediating stress associated with chromosome amplification has remained elusive. Ssd1 has been linked to translational regulation (including but not exclusively translational repression), mRNA localization, and PBs (J. M. Jansen et al. 2009; Ohyama et al. 2010; Kurischko, Kim, et al. 2011; Wanless, Lin, and Weiss 2014). It is known to impact cell wall biology, aging, and quiescence among other phenotypes (Kaeberlein and Guarente 2002; Xu et al. 2019; Miles et al. 2019). The apparent complexity of Ssd1’s role is not unique: many RBPs are pleiotropic, either in function (*e.g.* multiple roles within the cell) or in the phenotypes they impact. For instance, it is common for RBPs to participate in multiple, different stages of mRNA life cycles, or even to have secondary functions unrelated to their RNA-binding activity (reviewed in Hentze et al. 2018). Consequently, pinpointing the specific functions of Ssd1 that contribute to aneuploidy tolerance presents a significant challenge.

This study took a network approach to identify RBPs whose mRNA targets are shared with Ssd1, under the hypothesis that they may function together to regulate those mRNAs and impact aneuploidy tolerance. We found that Ssd1 shares more targets than expected with 14 other RBPs. Remarkably, 10 of these 14 RBPs (over 70%) influenced growth of cells with an extra copy of Chr12, when the RBP as deleted in wild-type aneuploids or in the context of *SSD1* deletion. We identified several RBPs whose deletion that sensitized cells to Chr12 duplication in their own right, albeit with much less effect that *SSD1* deletion, including Dhh1, Pat1, and to a lesser extent Mpt5 and others. Thus, this work identified new players in the modulation of Chr12 aneuploidy tolerance, and identified a refined network that suggests associated functions for future investigation.

Combining experimental results from several assays, interrogating Chr12 duplication but also others, presented a subnetwork of proteins that influence aneuploidy tolerance in *ssd1*Δ cells. One of the most striking commonalities among these proteins are their connections to translational repression and/or PB function. Poly-A binding protein Pab1, whose duplication benefits multiple *ssd1*Δ aneuploids (Dutcher et al. 2024), is important for efficient translation initiation but also plays a role in regulating repression and can also localize to PB (Chritton and Wickens 2010; Brambilla et al. 2019; Brengues and Parker 2007). The helicase Dhh1 and decapping factor Pat1 are also important for translational repression and are essential PB components: cells lacking both proteins have pronounced defects in PB formation and translational suppression (Coller and Parker 2005; Vijjamarri et al. 2023; Marnef and Standart 2010; Vindry et al. 2019; Zeidan et al. 2018). We found that deletion of either factor sensitizes cells to Chr12 duplication, even in the presence of Ssd1. Deletion of another PB factor, *MPT5*, also produced a mild sensitivity to Chr12 aneuploidy that was severe in the absence of *SSD1*. Mpt5 recruits specific mRNAs to PB, impacting either their decay or storage, depending on the target (Wang et al. 2018). Additional PB proteins were implicated by our network analysis (Figure 3B), including decapping and PB factor Lsm4, translational regulator Stm1 (which associates with nontranslating ribosomes, can promote decapping, and is a suppressor of *pat1Δ* temperature sensitivity) (Balagopal and Parker 2009; 2011), ribosomal protein Rps28b whose 3’ UTR serves as a scaffold for PB assembly (Fernandes and Buchan 2020), and translational repressor Sbp1 that also has a role in PB disassembly (Roy et al. 2022). Thus, the majority of proteins represented in the sub-network shown in Figure 3B are directly linked to PB physiology. The implication of this result is not clear, since the exact function of PB remains under investigation (reviewed in Escalante and Gasch 2021). PB have long been associated with translational silencing, although their formation may be a consequence, rather than a driver, of translational repression (Eulalio et al. 2007). Many PB components function outside of PBs in roles related to translational repression and/or mRNA decay. We hypothesize that their implication here underscores the importance of translational repression for aneuploidy tolerance and a specific role for Ssd1 in that process.

In contrast, several genes whose deletion ameliorates *ssd1Δ* aneuploidy sensitivity are associated with mRNA stability, especially at specific classes of mRNAs including cell-wall transcripts and mRNAs with upstream open reading frames (uORFS), both of which are enriched among Ssd1 targets. Nab6 stabilizes mRNAs encoding cell wall proteins (Bresson et al. 2023), while Pub1 stabilizes uORF-containing mRNAs by preventing nonsense-mediated decay (NMD) (Ruiz-Echevarría and Peltz 2000). Deletion of these RBPs is thus predicted to destabilize bound transcripts, which could counteract a loss of translational repression in the *ssd1*Δ strain. Interestingly, Ssd1 targets are enriched for both cell wall mRNAs (Hogan et al. 2008; Hose et al. 2020) and transcripts with uORFs (p = 9.4E-03, see Methods) (Ingolia et al. 2009; Spealman et al. 2023). Several genes encoding cell wall proteins (but also many others) are especially deleterious in *ssd1*Δ euploid cells and are encoded on Chr12, perhaps contributing to its toxicity (Dutcher et al. 2024). Another gene whose deletion alleviated the *ssd1Δ* Chr12-aneuploid growth defect was Ccr4, a core component of the Ccr4-Not deadenylase complex that influences mRNA decay, including of some cell wall mRNAs (Tucker et al. 2001; Ito et al. 2011).

Integrating these details suggests several models through which Ssd1 could function. Our results point to a role for Ssd1 in translational repression to mediate aneuploidy tolerance, of Chr12 studied here and other chromosomes investigated elsewhere (Dutcher et al. 2024; Rojas et al. 2024; Hose et al. 2020). We previously ruled out a model in which Ssd1 globally silences all amplified mRNAs encoded on the extra chromosome (Hose et al. 2020). Past proteomic work identified few proteins whose abundance increased in the *ssd1Δ* Chr12 aneuploid compared to the wild type aneuploid in a way that could not be explained by underlying mRNA changes; only 8 of the affected proteins come from Ssd1 targets, and only one of those is encoded on the amplified chromosome. Most Ssd1-bound transcripts are not more toxic when duplicated in isolation in *ssd1Δ* euploids, indicating that loss of repression of single transcripts is unlikely to drive aneuploidy sensitivity of *ssd1*Δ cells (Dutcher et al. 2024).

Instead, Ssd1 may help to mitigate indirect effects of aneuploidy on translational efficiency or fidelity. If chromosome amplification taxes translation, perhaps through over-abundance of many translated mRNAs, this could produce a cellular state that is sensitized to further translational stress. Several lines of evidence support a role for Ssd1 in managing that stress. First, aneuploid yeast are sensitive to the aminoglycoside nourseothricin (NTC), which binds the ribosome to disrupt elongation and cause tRNA misincorporation (Hose et al. 2020; Dutcher et al. 2024; Ling et al. 2012; Haupt et al. 1978). Deletion of *SSD1* renders aneuploid cells—but not euploids—extremely sensitive to NTC (Dutcher et al. 2024). Thus, the combination of *SSD1* deletion and chromosome amplification exacerbates sensitivity to this translational inhibitor. Second, *SSD1* deletion sensitizes euploid cells to mutation of the Elongator tRNA-modification complex. Loss of Elongator function increases tRNA misincorporation at specific codons (Karlsborn et al. 2014; Xu et al. 2019; 2020). Third, this connection between Ssd1 and translational fidelity may explain a recent result from other work in our lab that showed that over-expression of tRNAs improves growth of multiple *ssd1Δ* aneuploid strains (Rojas et al. 2024). Increased abundance of specific tRNAs can enhance translation through cognate codons, which may explain why tRNA up-regulation alleviates growth defects in several *ssd1*Δ aneuploids (Kramer and Farabaugh 2007; Rojas et al. 2024; Rak et al. 2018; Frumkin et al. 2018). Finally, we recently found that *SSD1+* aneuploids in the YPS1009 strain background can tolerate extra chromosome during log-phase growth, but they have a major defect entering quiescence and maintaining normal lifespan—these premature aging defects are due in part to defects in the Ribosome Quality Control (RQC) pathway that responds to stalled ribosomes (Sitron and Brandman 2020; Escalante et al. 2024). Remarkably, simply over-expressing stoichiometrically limiting RQC subunits partly alleviates aneuploid defects (Escalante et al. 2024).

One possibility is that Ssd1, and potentially other factors identified here, regulate mRNAs that are either error prone or difficult to translate. If aneuploid cells are already taxed, perhaps if over-abundant mRNAs from the amplified chromosome titrate key translation factors, deletion of Ssd1 and misregulation of specific targets could exacerbate problems. Interestingly, Kershaw et al. recently identified seven RNA regulons based on shared RBP interactions. Ssd1-bound mRNAs are overrepresented (p = 5.4E-15, hypergeometric test) in a regulon characterized by long, relatively structured mRNAs with low ribosome occupancy under standard conditions (Kershaw et al. 2023, “Cluster 3”). Indeed, Ssd1 targets are more structured relative to all yeast mRNAs (Kertesz et al. 2010) (p = 2.0E-06, see Methods). Thus, Ssd1 targets could have unique requirements for efficient translation, such that proper regulation of these mRNAs (or a subset thereof) is particularly important in aneuploids. Further studies will be required to elucidate the mechanistic underpinnings of these interactions, as well as to better understand the fundamental liabilities associated with aneuploidy stress. In all, this study identified new RBPs that influence aneuploidy tolerance, uncover novel genetic interactions with *SSD1* in the context of aneuploidy, and implicate functions important for tolerance of chromosome duplication.

## Methods

### Strains and growth conditions

Strains used in this study are listed in Table S1. Gene knockouts were generated by homologous recombination of HYG-MX into the designated locus, followed by diagnostic PCR to confirm absence of one copy of the target RBP gene. Colony based growth was scored as described below. Liquid growth curves were performed in YPD (1% yeast extract, 2% peptone, 2% dextrose) in test tubes at 30C with shaking. Specifically, liquid cultures were inoculated, grown for 3-4 hours to minimize extra-chromosome loss, and optical density at 600 nm (OD_600_) was measured over time to calculate exponential growth rates. Genetic interactions were determined as described below.

### mRNA target overlap analysis

We identified 65 proteins that shared targets with our 286 previously identified Ssd1-bound mRNAs (Hose et al. 2020) using the BioGRID database (Oughtred et al. 2016) accessed via Yeastmine (data obtained Oct. 2019) (Balakrishnan et al. 2012) along with several other direct studies available at the time (Delaveau et al. 2016; Shahbabian et al. 2014; Hogan et al. 2008). Of these 65 RPBs, 39 shared at least 3 RNA targets with Ssd1, where SSd1 targets were defined as those measured in our lab previously (Hose et al. 2020). We identified the subset of these RBPs that bound more Ssd1 targets than expected by chance, using hypergeometric tests (assessing a total of 5944 mRNAs) with Benjamini-Hochberg FDR correction (Benjamini and Hochberg 1995), taking FDR < 0.05 as significant. This identified the 14 RBPs shown in Figure 1.

### Genetic interaction analysis

A single copy of each RBP was deleted from diploid HO+ parental strain YPS1009 that was hemizygous for *SSD1* and harbored three copies of chromosome 12. The resulting double hemizygous strains (SSD1/ssd1*Δ*::KanMX; RBP/rbp*Δ*::HygMX) were sporulated and dissected to produce evenly spaced colonies of all desired genotypes (average of 22 colonies per genotype). Cells were grown on YPD plates for 3 days at 30C, imaged, scaled by 200%, and colony size was quantified using the R package gitter v 1.1 (Wagih and Parts 2014) using a reference image (see R package documentation) and plate.format=c(4,8), with otherwise default parameters. Colony genotype was subsequently determined based on segregation of drug markers (G418 resistance indicating *ssd1Δ::KANMX* or hygromycin resistance indicating *rbpΔ::HYGMX*). Flow cytometry was used to assess aneuploidy (described below). Tetrads with missing colonies or aberrant segregation patterns were excluded from downstream analysis.

The presence of synthetic genetic interactions was assessed as follows: for each independent dissection batch, the distribution of colony sizes was determined for each genotype. Genotype means were used to calculate defects for single mutants and aneuploids relative to wild-type controls. These defects were then used to calculate the distribution of expected colony sizes using a combinatorial model. For example, to generate the distribution of expected colony sizes for a given RBP mutant aneuploid, we first defined the defect due to Chr12 duplication using the mean size of the wild-type Chr12 colonies versus the mean size of wild-type euploid colonies, for colonies measured in the same batch. We then multiplied this defect by the observed size of each RBP mutant in the absence of aneuploidy, again for colonies in the same batch. The results generated a distribution of expected colony sizes that represent the expected combinatorial effect of aneuploidy and of RBP deletion in a euploid background. We then identified a statistically significant genetic interaction if the observed *rbpΔ* aneuploid colony sizes were greater, or smaller, than the expected colony sizes. A similar procedure was performed for other comparisons, estimating expected growth defects for each *ssd1Δ rbpΔ* euploid (*ssd1Δ* euploid defect x *rbpΔ* euploid defect) and *ssd1Δ rbpΔ* aneuploid (*ssd1Δ* aneuploid defect x *rbpΔ* aneuploid defect). Observed vs. expected colony sizes were compared using two-sided independent t-test with Benjamini-Hochberg FDR correction (Benjamini and Hochberg 1995).

Synthetic genetic interactions were assessed from liquid growth rates using a similar procedure. Growth rates were measured in at least biological triplicate for each strain, with strains being compared grown side-by-side in each replicate to enable replicate-paired t-tests to assess statistical significance.

### Flow cytometry to assess aneuploidy

To determine euploid/aneuploid status, a portion of each colony was scraped, resuspended in 70% EtOH and stored at 4 degrees. To process for flow cytometry, cells were incubated in 50ug/mL RNase A in 0.05 M sodium citrate for at least 3 hours or overnight, then incubated with 50ug/mL proteinase K in 50 mM TRIS pH 8.0 and 10mM CaCl2 for at least 3 hours or overnight. Processed cells were stained with 1uM Sytox Green for 15 minutes at room temperature, then run at a medium flow rate on Guava EasyCyte flow cytometer (Millipore). We defined differences in Sytox Green signal for known euploid and aneuploid control colonies. Aneuploidy genotyping was then done using these guidelines to assess all colonies of a given genotype from the same plate.

### Protein-protein interaction network and enrichments

Interaction data were obtained from BioGRID via Yeastmine as above and visualized using Cytoscape v. 3.9.1 (Shannon et al. 2003). Beneficial gene duplications were taken from (Dutcher et al. 2024), restricted to the 316 genes that were especially beneficial in Chr12 aneuploids. We identified the subset of genes related to RNA functions, starting with genes mapped to the following GO SLIM process terms: mRNA processing, rRNA processing, RNA splicing, regulation of translation, cytoplasmic translation, translational elongation, RNA modification, sno(s)RNA processing, tRNA processing, mitochondrial translation, and tRNA aminoacylation for protein translation, according to the Saccharomyces Gene Database (Cherry et al. 2012). We then manually curated annotations and added genes with related functions that were missing from these categories, then simplified categorization for visualization purposes. This process yielded the 67 genes depicted in Figure 3A (peripheral nodes plus *DHH1*). All genes shown in Figure 3A, including the RBPs with significant genetic interactions (central nodes), were then analyzed to determine if they had more interactions with central nodes than expected by chance as follows: first, we identified the subset of encoded proteins that interacted with at least 2 central node RBPs (see Figure 3A). For each protein that met this criterion, overrepresentation of interactions was then assessed using a hypergeometric test in R: phyper(x, m, n, k, lower.tail = FALSE), where x = the number of central node interactions for the given protein; m = the total number of interactors for that protein; n = 6400 (estimated size of total pool of interactors) – m; and k = 11 (number of central RBP nodes). Benjamini-Hochberg correction was then used to calculate FDR, taking FDRs < 0.05 as significant. Functional enrichments were assessed using the hypergeometric function and GO annotations defined in setRank version 1.0 (Simillion et al. 2017). Genes shown in network figures were manually categorized according to their primary functions, as listed in the key. Of the 286 Ssd1 RNA targets identified in Hose et al. 2020, 57 transcripts with uORFs were identified using data from Ingolia et al. 2009, combining a list of predicted uORFs with translated uORFs identified in that study. Analysis of secondary structure of Ssd1-bound RNAs compared to the whole transcriptome was conducted using numerical scores representing secondary structure from Kertesz et al. 2010. Distributions of all available scores corresponding to the coding regions (CDS) of Ssd1 bound mRNAs from Hose et al. 2020 were compared to total available CDS scores the yeast transcriptome using a two-sided Mann Whitney U test. Ssd1 target overlap with clusters identified in Kershaw et al. 2023 were determined using hypergeometric test: 96 of 267 Ssd1 targets (Hose et al. 2020) present in the dataset are in “Cluster 3,” out of a total of 5050 RNAs across all clusters (Kershaw et al. 2023).

## Acknowledgements

This work was funded by NIH grants R01CA229532 and R01GM148975 to APG. HAD was supported by training grants T32GM007133 and T32HG002760 to the Genomic Sciences Training Program.

